# Neural Correlates of Task-related Refixation Behaviour

**DOI:** 10.1101/773143

**Authors:** Radha Nila Meghanathan, Cees van Leeuwen, Marcello Giannini, Andrey R. Nikolaev

**Author notes:** Corresponding author Center for Cognitive Science, Technical University Kaiserslautern, 57 Erwin-Schrӧdinger-Strasse, Kaiserslautern 67663, Germany.

## Abstract

Information uptake during scene viewing under free viewing conditions is crucially determined by the scanning plan. This plan is determined both by top-down and bottom-up factors. To capture top-down factors affecting saccade planning, we compared EEG between first fixations and refixations on items varying in task-relevance. First fixations and refixations impose different working memory costs because first fixations involve encoding of new items whereas refixations involve rehearsal of existing items in working memory. These memory requirements also differ with the task-relevance of the item being encoded. Together, these two factors of task-relevance and memory processes related to refixation behavior would affect saccade planning. In a visual task involving search and memorization of multiple targets, we compared saccade-related potentials (SRPs) between first fixations and refixations for task-relevant (target) and task-irrelevant (distractor) items. We assessed the interval preceding a saccade away from the fixation of interest. Studying this presaccadic interval revealed how mechanisms related to saccade preparation are affected by task-relevance and refixation behavior. We found higher SRP amplitudes for first fixations than refixations over the occipital region for task-relevant items only. Our findings indicate that saccade planning is modulated by both task-relevance of an item and working memory load.

## 1. Introduction

Before a saccade is executed, attention shifts to the next fixation location (Deubel & Schneider, 1996) after *disengagement* of attention from the currently attended location (Posner & Petersen, 1990). Attention shift is biased by the current contents and load of visual working memory (Van der Stigchel & Hollingworth 2018). This bias is seen in free viewing conditions where fixations are guided not only by bottom-up factors such as salience and spatial layout but also by top-down factors such as task and prior knowledge (Baluch & Itti, 2011; Henderson, 2003, 2007; Rayner, 2009; Tatler, Hayhoe, Land, & Ballard, 2011). We will focus on top-down factors that contribute to saccade planning under free viewing conditions.

Top-down factors may play different roles when the eyes first fixate an object or when they return to a previously fixated one (Yarbus, 1967). Eye-tracking research typically assigns to such *refixations* the role of updating (Tatler, Gilchrist, & Land, 2005) or rehearsing (Meghanathan, Nikolaev, & van Leeuwen, 2019; Zelinsky, Loschky, & Dickinson, 2011) object representations in visual working memory (VWM). Object representations need updating or rehearsing in the case of insufficient initial examination (Beck, 2006; Gilchrist & Harvey, 2000; Peterson, Beck, & Wong, 2008; Peterson, Kramer, Wang, Irwin, & McCarley, 2001), poor or decayed memory (Körner & Gilchrist, 2008; Meghanathan et al., 2019; Zelinsky et al., 2011) or for acquisition of task-specific information (Droll, Hayhoe, Triesch, & Sullivan, 2005; Hayhoe, Bensinger, & Ballard, 1998). This suggests that initial examination of an item (first fixation) and refixation are distinct in executive and working memory processes. First fixations involve encoding into VWM, whereas refixations involve updating or reactivation of a previously encoded item. Since first fixations alone involve memorization of the item at fixation, disengagement of attention may be more difficult compared to that in refixations. Accordingly, we may expect a difference in saccade planning after first fixations and after refixations.

However, this difference is likely to be modulated by the task-relevance of an item, that is, whether it is a target or a distractor. The difference in costs of disengagement between the first fixation and refixation will, accordingly, be relevant for targets, processing of which requires memorization of both identity and location. Since identity of distractors is not task-relevant, refixations to distractors are likely made because of forgetting of distractor locations (Meghanathan et al., 2019). Consequently, first fixations and refixations on distractors should involve similar memory uptake. Disengagement of attention from the currently fixated distractor should not be different between first fixations and refixations. Hence, for distractors, saccade planning during a refixation is unlikely to be different from that during first fixation.

We propose that saccade planning in refixations differs from that of first fixations for targets but not for distractors. To test this hypothesis, we analyzed EEG in the interval of fixation on targets and distractors in an experiment involving a multitarget visual search task. The results of the refixation analysis in this experiment were recently published elsewhere (Meghanathan et al., 2019). We observed that refixation behavior depends on both task-relevance of the refixated item and on working memory load. Here, we focus on neural mechanisms of oculomotor selection, using the same dataset which includes EEG coregistered with eye movements. Specifically, we compare EEG between the first visit (*first fixations*) and subsequent visits (*refixations*) of task-relevant (targets) and task-irrelevant (distractors) items.

In free viewing, the fixation interval is likely to involve both visual processing of the currently fixated item (Henderson, 2007; Irwin, 2004) and covert attention to the subsequent item to be fixated (Kovalenko & Busch, 2016; Kowler, Anderson, Dosher, & Blaser, 1995). Visual processing and saccade preparation can be distinguished by time-locking of the EEG activity to the onset of the current fixation and the following saccade, respectively (Nikolaev et al. 2016). Since our hypothesis was related to disengagement of attention from the current to the next fixation location, we measured EEG time-locked to the onset of the saccade following a target or distractor fixation. This presaccadic or saccade-related potential (SRP) has been known to indicate saccadic preparation (Gutteling, van Ettinger-Veenstra, Kenemans, & Neggers, 2009; Wauschkuhn et al., 1998), attention shift to saccade location (Gutteling et al., 2009; Kovalenko & Busch, 2016; Krebs, Boehler, Zhang, Schoenfeld, & Woldorff, 2011; Wauschkuhn et al., 1998) and transsaccadic remapping (Parks & Corballis, 2008, 2010). In an earlier study, we found a difference in presaccadic potentials, and therefore, saccade planning between ordinary fixations and refixations (Nikolaev, Meghanathan, & van Leeuwen, 2018). This indicated a difference in presaccadic attention shift between fixations and refixations.

## 2. Material and Methods

### 2.1 Participants

Twenty three participants (16 female) with mean age of 20.86 years (ranging between 18 and 29 years) took part in the experiment with informed consent. The study was approved by the Ethics Committee of KU Leuven. Data from two participants with excessive eye movements outside the monitor were excluded. Of the remaining 21 participants, 20 (15 female) met the eye movement criteria described later (Section 1.6). Their mean age was 21.75 years.

### 2.2 Stimuli and Procedure

In each trial, we presented a visual search display (Figure 1) followed by a change detection display, both subtending 39.9° × 30.5° of visual angle. There were 40 black items (0.48 cd/m^2^) in each display presented on a grey background (32.84 cd/m^2^) of which 3, 4, or 5 were target Ts (0.41° × 0.41°) and the rest distractor Ls (0.31° × 0.41°), randomly placed within a rectangular space of visual angle 32.9° × 23.12°. All items were separated by a minimum of 3.12°, while targets were separated by at least 6.24°. All items were encircled (0.41°) to lower the possibility of peripheral detection (Körner & Gilchrist, 2007; Peterson et al., 2001). Both targets and distractors were randomly oriented in one of 20°, 80°, 140°, 200°, 260° or 320° clockwise from the vertical. All targets in a display had different orientations. All items occurred in the same locations and in the same orientations in both displays except for one target, which was oriented differently in the change detection display in one half of the trials.

**Figure 1.**
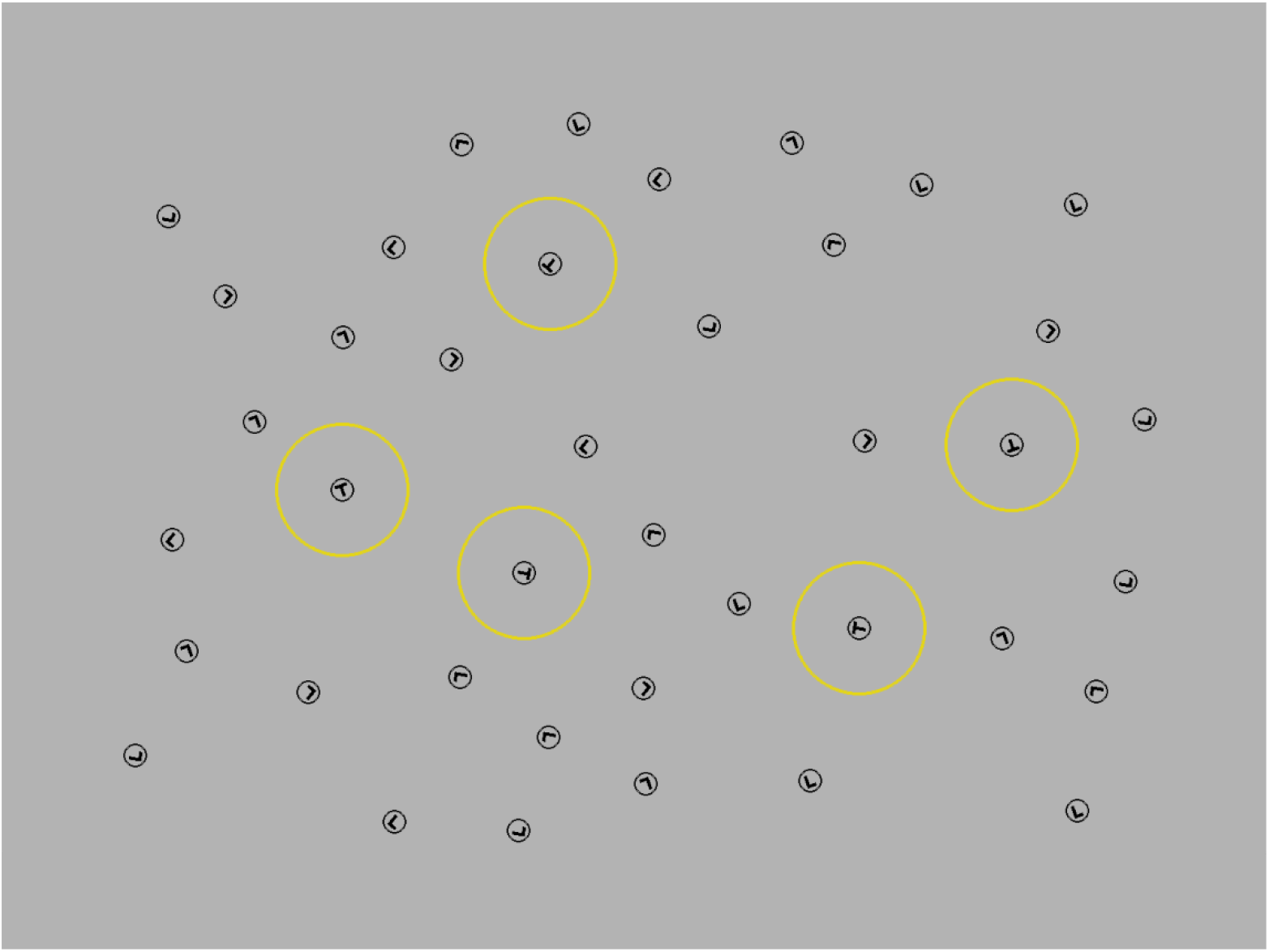
A sample display with 5 targets and 35 distractors. Targets are highlighted in yellow circles of 2° radius.

Participants fixated on a central fixation dot and initiated each trial by pressing a key. First, the visual search display was presented for 10 s, during which time participants were asked to search for 3, 4 or 5 targets and memorize their orientations. Participants were not told how many targets would occur in each trial. After an interval that randomly lay between 1 and 1.5 s, the change detection display was presented until participants indicated whether they detected a change or not by pressing one of two keys within 10s. After this, a feedback display was presented with the targets in green circles in the case of a correct response, targets in red circles in the case of an incorrect response and with the changed target in a large yellow circle in trials where target orientation was changed.

### 2.3 Experiment setup

There were 270 trials in the experiment with 90 trials in each target condition (3, 4 or 5). Participants performed trials in 10 blocks of 27 trials each with at least a two-minute break between the blocks. Target conditions were randomized across the 10 blocks. The experiment lasted about an hour and forty minutes.

The experiment setup involved a presentation computer, an EEG recording computer and an eye movement recording computer. The stimuli were presented by the presentation computer on a 30 cm × 40 cm monitor with a screen resolution of 1600 × 1200 and a refresh rate of 75 Hz. Participants were seated 55 cm from the monitor with their head stabilized on a chin rest. Trial onset and offset events were synchronized between the EEG and eye tracking systems by a TTL signal sent via a parallel port from the presentation computer.

### 2.4 Eye movement recording

Eye movement data was recorded using an EyeLink 1000 eye tracker on a desktop mount (SR Research Pvt. Ltd., Ontario, Canada) at a sampling frequency of 250 Hz. A 9-point calibration was performed at the start of the experiment for the left eye with calibration points at 4 corners and 4 mid-points of the edges of the display. If error was less than 2°, calibration was considered successful and the left eye was tracked during the experiment. If calibration was consistently poor for the left eye, the right eye was calibrated and tracked. The eye was calibrated before the start of each block. Additionally, drift of the eye was corrected before the start of each trial when participants fixated on the central fixation dot. If error due to drift exceeded 2°, calibration was repeated.

### 2.5 EEG recording

EEG was recorded using a 256 channel Geodesic Sensor Net (EGI, a Philips company, Eugene, OR) at a sampling frequency of 250 Hz with Cz as the reference electrode. Sensors for recording horizontal and vertical EOGs were part of the electrode montage. During recording, the input signal was filtered with analog filters at a low cut-off of 0.1 Hz and a high cut-off of 100 Hz.

### 2.6 Eye movement preprocessing

Eye events detected by the EyeLink software were used in the analysis. When eye velocity exceeded 22°/s and acceleration exceeded 3800°/s^2^, a saccade was detected. The intervals between saccades were categorized as fixations.

#### 2.6.1 Preprocessing of fixation sequences

We analyzed data from the visual search stage of the task for all trials with accurate responses. The first fixation in each trial was excluded because of overlap of fixation-related activity and EEG evoked by the screen onset (Dimigen, Sommer, Hohlfeld, Jacobs, & Kliegl, 2011). We included fixations of duration less than or equal to 1000 ms in the analysis, which was 99.9% of the total number of fixations. Fixations were assigned either to a target or a distractor when the fixation was within 2° from the center of the item. In line with previous studies (McCarley et al., 2006; McCarley, Wang, Kramer, Irwin, & Peterson, 2003; Peterson et al., 2001), a 2° criterion ensured that the fixated item lay within the fovea. 8% of fixations on average were assigned to neither target nor distractor and were not included in the current analysis.

#### 2.6.2 Selection of first fixations and refixations

From the fixation sequence of each trial, fixations that were first visits to targets and distractors were identified. These were called first fixations. After this first visit, any fixation on the same item was considered a refixation. Note, however, that the first fixations were considered independently on whether or not the item was revisited. We analyzed EEG time-locked to the onset of the saccade following the current fixation (Figure 2).

**Figure 2.**
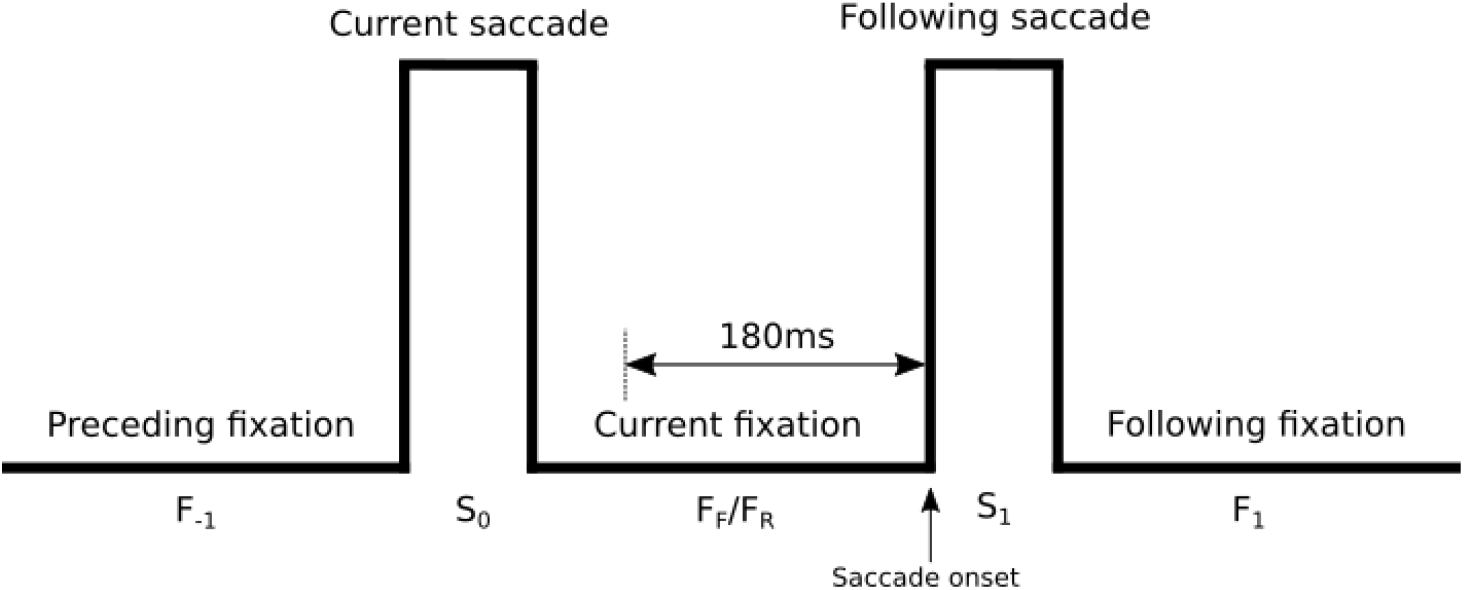
Schematic depicting first fixation, F_R_ or refixation F_R_ on a target or distractor. Saccade onset indicates the onset of the saccade following a fixation or refixation on a target or distractor. EEG was segmented relative to the saccade onset, S_1_, for analysis of the eventual 180ms of a fixation or refixation.

We ensured that the saccade preceding and following the fixation of interest was always made to a distractor to rule out any item-based bias on covert attention in the presaccadic interval. Since the type of the following item was the same across conditions, any difference between the fixation types would be specific to the task-relevance (target or distractor) of the currently fixated item. To achieve this, we applied exclusion criteria on the candidate fixations for the first fixation condition and refixation condition for EEG analysis based on the preceding and following fixations as depicted in Figure 2. Let F_F_ be the current first fixation and F_R_ be the current refixation being considered. F_R_ was not included in the analysis if F_−1_ or F_1_ was on the same item. F_F_ was not included in the analysis if F_1_ was on the same item. Additionally, F_F_ or F_R_ was excluded if F_1_ was on a target.

After identification of fixations for the first fixation and refixation conditions, we applied threshold criteria for inclusion of fixations. We removed current saccades, S_0_, shorter than 20 ms and longer than 80 ms. In addition to exclusion of current fixations longer than 1000 ms (see above), we also removed fixations, F_F_/F_R_, shorter than a criterion of 180 ms. It was imperative that the fixation duration of F_F_ or F_R_ be equal to or greater than the length of an EEG epoch, since otherwise an EEG epoch may include intervening saccades which distort EEG. The threshold of 180 ms was selected as a trade-off between maximizing both EEG epoch durations and number of participants with the criterion number of epochs. This criterion was set at 40 epochs per condition to include a participant in the EEG analysis.

##### 2.6.2.1 Matching of eye movement characteristics

In a free viewing task, EEG response to cognitive processes is confounded by that to eye movements (Dias, Sajda, Dmochowski, & Parra, 2013; Dimigen et al., 2011; Nikolaev, Meghanathan, & van Leeuwen, 2016). For instance, we have shown that saccade size and fixation duration affect EEG on the current fixation (Nikolaev et al., 2016). Therefore, we need to ensure that the effect of experimental conditions on EEG are not due to systematic differences in eye movements between the conditions. This can be done either by finding eye movements with matched characteristics across experimental conditions (Dias et al., 2013; Nikolaev et al., 2016) or by including eye movement characteristics as predictors in a regression model (Cornelissen, Sassenhagen, & Võ, 2019; Dandekar, Privitera, Carney, & Klein, 2012; Dimigen & Ehinger, 2019; Ehinger & Dimigen, 2018; Van Humbeeck, Meghanathan, Wagemans, Leeuwen, & Nikolaev, 2018). Both approaches have their pros and cons. In selecting eye events with matched eye movement characteristics, we are forced to exclude parts of the dataset. However, this approach allows us to interpret the results with reference to traditional EEG. Use of regression models, on the other hand, renders complex output often owing to interactions between predictors making interpretation of the results difficult. However, it also allows us to use the entire dataset for analysis.

For the current EEG analysis, we decided to use matching of eye movement characteristics. We followed guidelines for the matching procedure that we expounded in earlier work (Nikolaev et al., 2016). Since, we analyzed EEG in the concluding interval of a fixation, we matched the saccade duration (S_0_ in Figure 2) and fixation duration (F_F_/F_R_) of the candidate fixation between conditions (first fixations and refixations). In addition, we matched the rank of fixations between first fixation and refixation conditions to ensure that any difference we find is not due to their natural successive order in the course of a trial, which is known to affect fixation-related EEG (Fischer et al., 2013; Kamienkowski et al., 2018; Nikolaev et al., 2018). We matched epochs in the conditions of first fixations and refixations separately for targets and distractors. We matched first fixation and refixation epochs for each parameter (saccade duration, fixation duration and rank) sequentially by removing epochs till the conditions were matched as explained below. For each participant, we first arranged all refixation epochs by increasing order of saccade duration. Iteratively, we removed a refixation epoch and compared saccade duration between first fixation and refixation conditions via a *t*-test (*α* = 0.9), until a match was achieved. If fixations in the refixation condition had larger mean saccade durations than those in the first fixation condition for the participant, we removed refixation epochs from the higher end of the sorted saccade durations. In the opposite case, we removed refixation epochs from the lower end of the sorted saccade durations. Refixation epochs were removed at this step because of their larger number in comparison to first fixation epochs. Using the matched list of epochs of refixations for saccade durations, we repeated the same matching procedure for fixation durations. After this step, the number of epochs in refixation condition had reduced drastically. Therefore, in the next step of matching ranks, we repeated the same matching procedure but iteratively removed epochs of first fixations instead of refixations.

After matching within a participant, we checked if the two conditions were matched at the group-level. To that end, we tested the difference in means across participants via a paired *t*-test. Here we selected a very high *α* criterion of 0.9 for matching within a participant because it ensured a match at the group level. While saccade duration and fixation duration were matched, rank was not matched across participants even though it was matched individually for each participant. Therefore, we additionally matched ranks between targets and distractors for first fixation and refixation conditions separately to ensure that any EEG difference due to the difference in rank between the first fixation and refixation conditions is equally inherent in target and distractor conditions. Since the number of distractors was larger than that of targets, for each participant, we iteratively removed epochs in the distractor condition until a match was obtained using the procedure described above. After matching, the number of first fixation and refixation epochs on distractors was still disproportionately larger (about 3 times in some cases) than that on targets. Therefore, from the rank-matched distractor epochs, we randomly selected a subset of epochs of first fixations and refixations equal in number to the epochs of target first fixations and refixations for each participant. After this step, we checked for group-level match of (paired *t*-test, *α* = 0.05) saccade duration and fixation duration between distractor first fixations and refixations, and group-level match of rank between targets and distractors separately for first fixations and refixations. The random selection of distractor epochs was repeated until a match was obtained at the group-level. This gave us the final set of epochs of distractor first fixations and refixations that were used in the EEG analysis. The mean saccade durations and fixation durations for first fixation and refixation conditions of targets and distractors are in Table 1. For a fixation duration threshold of 180 ms, the matching procedure gave us 20 participants with at least 40 epochs in each condition (mean number of epochs = 160, SD = 30).

**Table 1.** Mean and SD of saccade and fixation durations for target and distractor fixations and refixations for 21 participants before and after applying limits on eye movement characteristics, and for 20 participants after matching.

|  | Target |  | Distractor |  |
| --- | --- | --- | --- | --- |
|  | Saccade duration (in ms) | Fixation duration (in ms) | Saccade duration (in ms) | Fixation duration (in ms) |
| Before applying limits |  |  |  |  |
| First fixation | 35.7 (4.5) | 279.4 (75.9) | 36.1 (4.3) | 195.9 (22.5) |
| Refixation | 40.9 (5.1) | 243.5 (34.9) | 37.3 (4.4) | 193.2 (20.6) |
| After applying limits |  |  |  |  |
| First fixation | 35 (3.2) | 317.3 (79.9) | 36.3 (3.2) | 230 (13.98) |
| Refixation | 40.2 (3) | 281.9 (34.5) | 36.7 (2.7) | 232 (11.7) |
| After matching |  |  |  |  |
| First fixation | 35.3 (3.4) | 323.1 (75.8) | 37.1 (3.5) | 234.3 (11.7) |
| Refixation | 35.1 (3.4) | 319 (80.4) | 36.6 (2.8) | 233.4 (11.4) |

### 2.7. EEG preprocessing

We processed EEG data using BrainVision Analyzer (Brain Products GmbH, Gilching, Germany). EEG data was first filtered with Butterworth IIR filters at a low cut-off of 0.5 Hz and a high cut-off of 30 Hz. We removed 95 electrodes close to the cheek and neck area with strong muscle and movement artifacts. We then visually inspected EEG channels to remove noisy ones. Data for these noisy channels were later interpolated from neighboring channels using spherical spline interpolation. The vertical electrooculogram, VEOG, was derived as the difference between electrodes placed above and below the eye, while the horizontal electrooculogram, HEOG, was derived as the difference between electrodes placed on the right and left outer canthi of the eye. After removing eye electrodes, we were left with 148 electrodes that we used in the EEG analysis.

For SRP analysis, EEG data was segmented in the interval −180 ms to 60 ms around the onset of the saccade that followed the fixation of interest. The interval of interest was the 180 ms preceding the saccade onset, which comprised of the eventual 180 ms of a first fixation or refixation.

We performed independent component analysis (ICA) to remove ocular artifacts. Data consisting of the concatenated segments was fed to an extended Infomax ICA algorithm. An ICA component was rejected when the sum of squared correlations with the HEOG or VEOG channel exceeded 0.3. Data was reconstructed from the remaining ICA components.

After ICA, the relevant epochs for EEG analysis were segmented in the interval −180 ms to 0 ms from the onset of the saccade following the fixation of interest. These segments were assessed for artifacts. An artifact was detected when EEG amplitude exceeded 50 μV between two subsequent samples, when amplitude difference exceeded 100 μV in an interval of 50ms or when activity was below 0.5 μV in an interval of 100 ms. Segments with artifacts were rejected.

### 2.8. EEG analysis

Since we had no specific predictions about the brain regions involved, we took two approaches to analyzing ERPs. In the first approach, we reduced dimensionality of our high-density electrode space by averaging across major regions of the brain. We defined 8 regions of interest (ROIs) corresponding to electrodes in the 10-20 system of placement of electrodes: frontal left and right (FL, FR), central left and right (CL, CR), parietal left and right (PL, PR) and occipital left and right (OL, OR). These ROIs were obtained by averaging one central electrode corresponding to the electrode in the 10-20 system (F3, F4, C3, C4, P3, P4, O3, O4) and the surrounding six electrodes.

We corrected each EEG segment using the interval −180 ms to −160 ms before the saccade onset as a baseline. To obtain the averaged SRP, we averaged EEG amplitudes across segments for each condition for each participant. We, then computed the mean amplitudes in the interval from −160 ms to 0 ms, which were then compared statistically between conditions. Separately for targets and distractors, we compared averaged EEG amplitudes of first fixations and refixations using 8 × 2 repeated measures ANOVAs with ROI (8 ROIs) and fixation type (first fixation, refixation) as factors. In case of violation of sphericity (Mauchly’s test), we used Huyhn-Feldt corrected *p*-values. Statistical analyses were done in R using the ez package (Lawrence, 2016).

The above analysis with 8 ROIs included only 56 of the 148 electrodes available. Besides, the comparisons were done on EEG amplitudes for the entire 160 ms interval. This may have ignored finer spatiotemporal differences between the fixation types. To complement the analysis with *a priori* selection of ROIs and time windows, in another approach, we compared baseline corrected data from all electrodes at all time points (160 ms) to locate the difference between the two fixation types in time and space. In order to do this, we performed the spatiotemporal cluster-based permutation test (Maris & Oostenveld, 2007), implemented in MNE-Python (Gramfort et al., 2014). Clusters were identified as contiguous points in space (electrode) and time (time point in EEG segment), which showed a significant difference between conditions via a *t*-test (*p* = 0.001). After adding *t*-scores of the samples in a cluster to obtain a cluster-level *t*-score, clusters were compared to a permutation distribution constructed from the largest cluster *t*-scores obtained in 1000 random assignments of data points to the two conditions. The rank of each cluster’s *t*-score in the permutation distribution was the probability, *p*, of finding a cluster of that *t*-score. Clusters of *p* larger than 0.001 were considered for comparing the two conditions. We expected this analysis to validate and corroborate the results from the *a priori* selection analysis.

## 3. Results

### 3.1. Eye movement results

The average current saccade duration and fixation duration for target and distractor fixations in the first fixation and refixation conditions are summarized in Table 1.

Before applying limits and matching, we compared saccade duration and fixation duration between first fixation and refixation conditions for targets and distractors. We used a repeated-measures ANOVA with the factors of task-relevance (targets vs. distractors) and fixation type (first fixation vs. refixation).

Saccade duration was longer for targets than for distractors (*F*(1,19) = 34.9, *p* < 0.001) and for refixation than first fixation (*F*(1,19) = 41.4, *p* < 0.001). But the interaction between task-relevance and fixation type (*F*(1,19) = 35.3, *p* < 0.001) indicated that there was no difference between targets and distractors for first fixation (post-hoc *p* = 0.48).

Fixation duration was longer for targets than for distractors (*F*(1,19) = 50.3, *p* < 0.001) and for first fixation than refixation (*F*(1,19) = 13.3, *p* = 0.002). But the interaction between task-relevance and fixation type (*F*(1,19) = 13.4, *p* = 0.002) indicated that there was no difference between first fixation and refixation for distractors (post-hoc *p* = 0.67).

### 3.2. EEG results

We compared EEG amplitudes during first fixations and refixations on targets and distractors separately. For targets, we found a significant effect of ROI (*F*(7, 133) = 5.3, *p* < 0.001, ε = 0.4) and of fixation type (*F*(1,19) = 12.99, *p* < 0.001) on EEG amplitudes (Figure 3). There was also an interaction between ROI and fixation type (*F*(7,133) = 14, *p* < 0.001, ε = 0.4). EEG amplitudes of refixations were greater than those of first fixations in FL and FR (both *p* < 0.05). The opposite was seen in PL, PR, OL and OR (all *p* < 0.05). For distractors (Figure 4), we found no significant effects.

**Figure 3.**
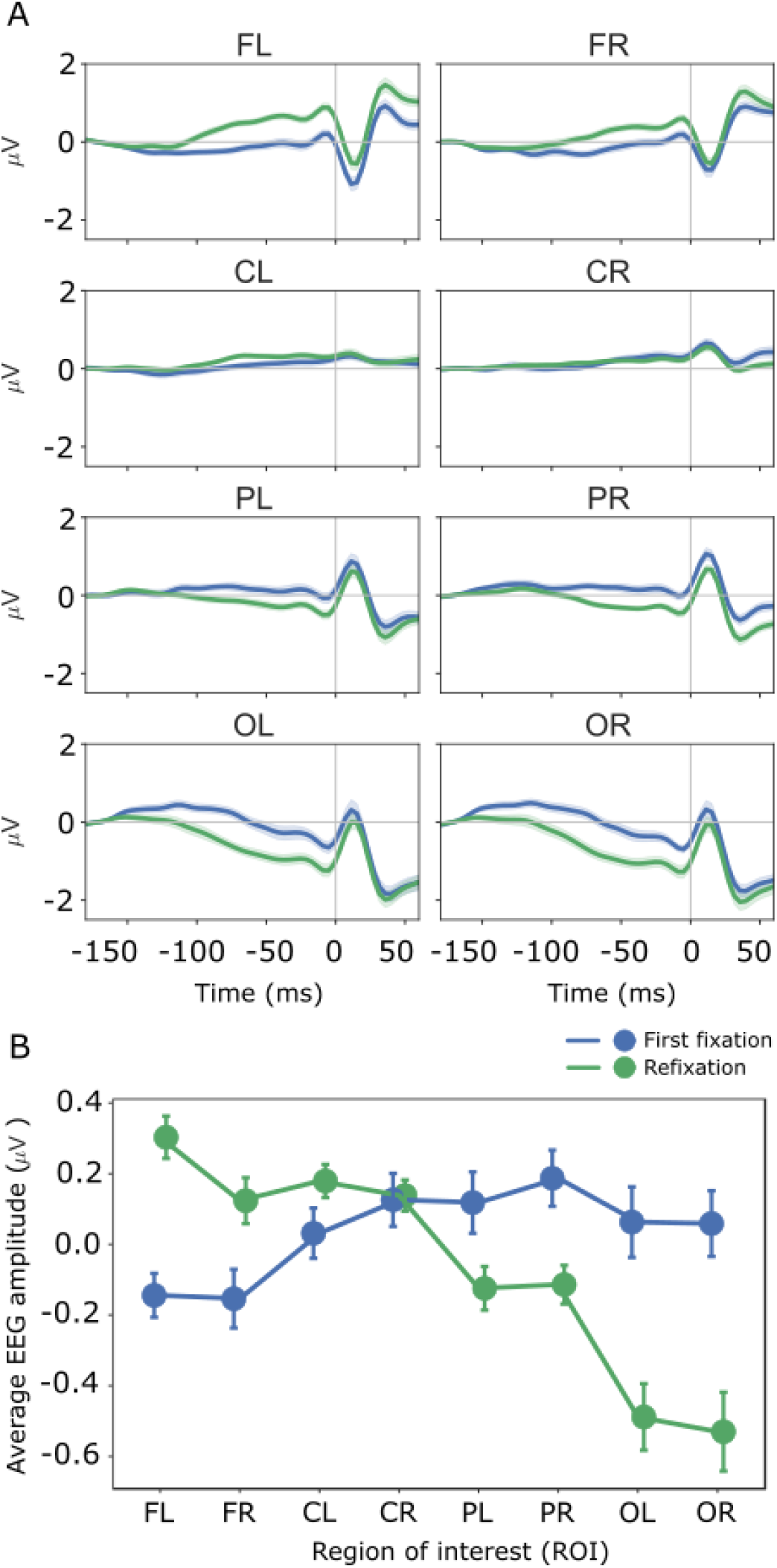
EEG amplitudes for targets. A. Grand-averaged potentials for 20 participants during first fixations and refixations on targets. 0 ms is saccade onset. The shaded area indicates 95% confidence intervals. B. The mean EEG amplitudes in the interval from −160 ms to 0 ms before the saccade onset for target first fixations and refixations for 20 participants. Error bars indicate standard error of mean.

**Figure 4.**
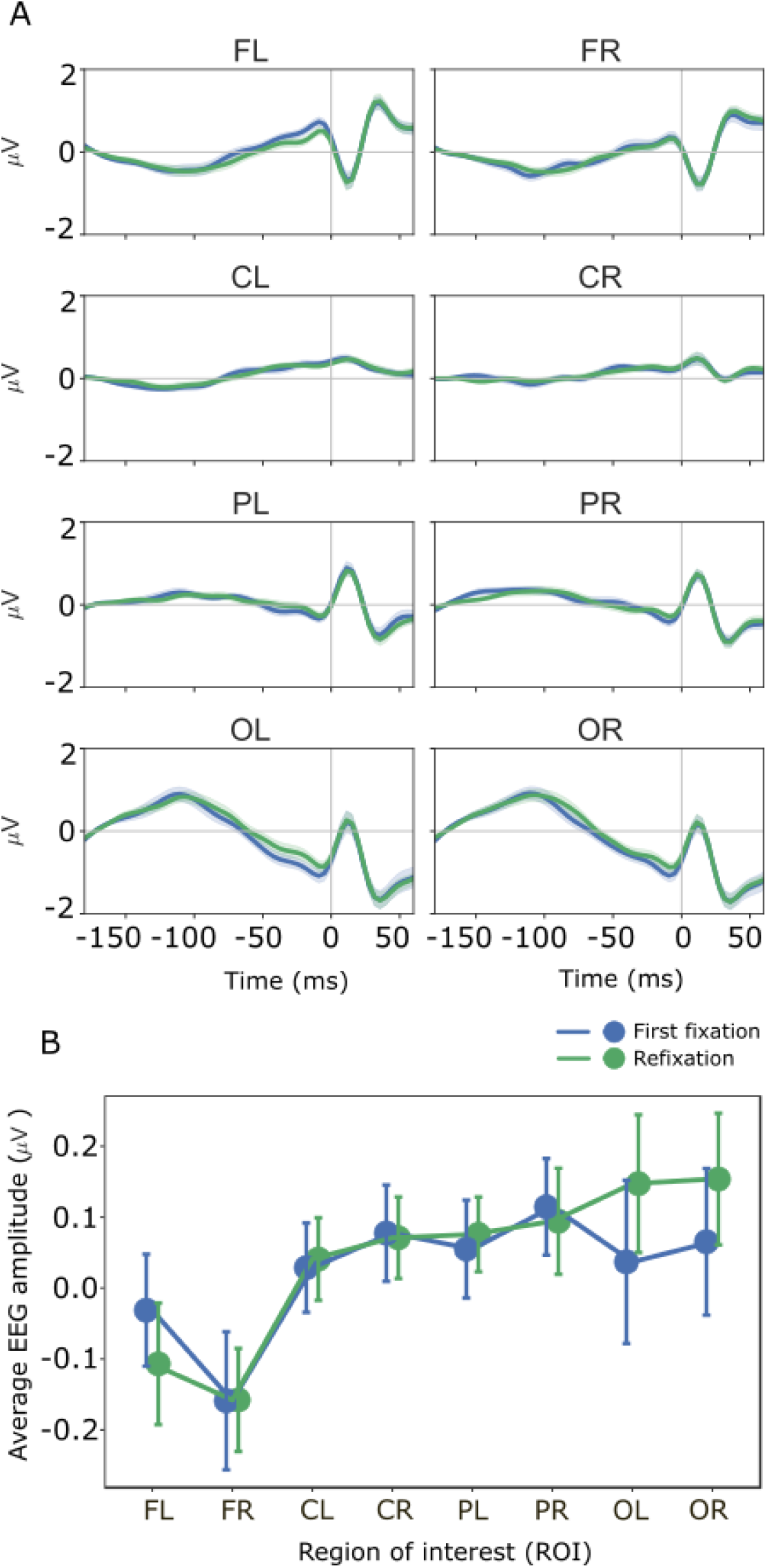
EEG amplitudes for distractors. A. Grand-averaged potentials for 20 participants during first fixations and refixations on targets. 0 ms is saccade onset. The shaded area indicates 95% confidence intervals. B. The mean EEG amplitudes in the interval from −160 ms to 0 ms before the saccade onset for distractor first fixations and refixations for 20 participants. Error bars indicate standard error of mean.

Using the cluster-based permutation test, we found two significant differences between first fixations and refixations for targets over occipital and frontal regions (Figure 5). The occipital cluster comprised 52 electrodes with a difference approximately from −112 ms to −8 ms. EEG amplitudes for first fixations were larger than that for refixations, similarly to the result of the ROI analysis. The frontal cluster included 30 electrodes primarily over the left frontal area differing approximately from −100 ms to −8 ms. As seen earlier for the frontal electrodes in the ROI analysis, refixations had larger EEG amplitudes than first fixations.

**Figure 5.**
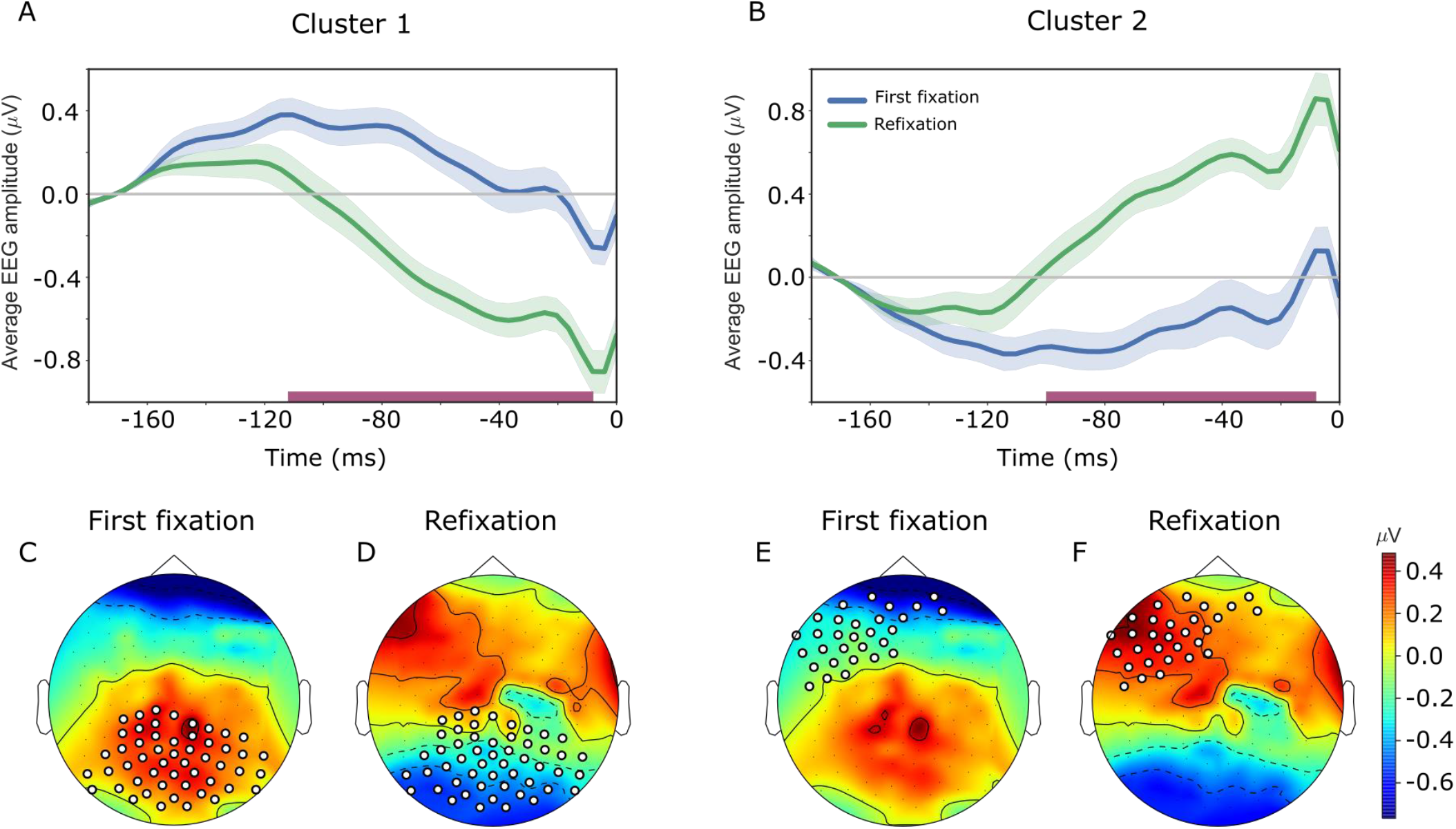
Results of the cluster-based permutation test on EEG amplitude for target first fixations and refixations. A. SRP averaged over 52 electrodes forming the occipital cluster. The pink bar along the x-axis indicates the interval of significant difference. B. SRP averaged over 30 electrodes forming the frontal cluster. 0 ms is a saccade onset. C-F: Topographic maps of EEG amplitudes averaged in the intervals indicated by the pink bars. C. The occipital cluster during first fixations. D. The occipital cluster during refixations. E. The frontal cluster during first fixations. F. The frontal cluster during refixations. The electrodes of the clusters are shown in white.

We did not find any difference between first fixations and refixations on distractors.

## 4. Discussion

Neural correlates of task-relevance during refixation behavior, to the best of our knowledge, have been never studied in humans. We raised the issue, whether preparation for a saccade following first fixations and refixations on an item depends on its task-relevance, i.e., whether it is a target or a distractor. To address this issue, we compared EEG between first fixations and refixations on targets and on distractors in a multitarget visual search task. We considered the saccade-related potential (SRP) in a 160-ms interval before the execution of the saccade following the fixation of interest. In general, visual processing of the fixated item and preparation for the subsequent saccade occur in this interval. But, since we used averaging relative to the onset of the following saccade in the EEG analysis, we can only consider processes of saccade planning and preparation rather than processes of information acquisition which are time-locked to the fixation onset.

We found a difference in the amplitude of SRPs between first fixations and refixations for targets. SRP amplitude was higher for first fixations than refixations over occipital areas with the opposite trend over the frontal regions (Fig. 3, Fig. 5). There was no difference in SRP amplitude for distractors (Fig. 4). This result indicates that preparation of the following saccade depends on task-relevance of the currently fixated item. We proposed that the difference between first fixations and refixations is likely caused by their different costs to VWM. During a first fixation on a target, attention is deployed to the target and it is encoded in VWM. Whereas, during a refixation, it is likely that an existing but deteriorating target representation in VWM is updated.

Consequently, the VWM processes during a refixation are different from those for a first fixation. The shift from one spatial location to another involves costly disengagement of attention (Posner & Peterson, 1990). Attention shift to the following saccade from the current fixation requires disengagement of attention, which is less taxing on VWM in the case of refixations than first fixations on targets. One reason for this may be that the VWM load during first fixations, which involves encoding, is higher than that during refixations, which involves only updating. This understanding is supported by the much longer duration (36 ms) of first fixations than refixations on targets before matching (Table 1). Since first fixations generally precede refixations, their durations would be expected to be shorter than refixations in keeping with the well-known increase in fixation duration during the course of a trial (Unema et al., 2005; Pannasch et al., 2008). The opposite effect in our data, instead, implies a difference in memory load, since fixation duration is a sensitive indicator of memory load (He & McCarley, 2010; Peterson et al., 2008, Meghanathan et al., 2015). Despite fixation duration being matched between first fixations and refixations for the EEG analysis, the VWM contents underlying EEG in the matched epochs may still differ in memory load. Thus, disengagement of attention may be more effortful for first fixations, when memory load is high, than for refixations, when memory load is low.

During fixations on distractors, VWM is not loaded with an extra item for maintenance since identities of distractors do not need to be remembered to perform the task. This is indicated by the very small difference (2 ms) of duration of first fixations and refixations before matching (Table 1). Only distractor location needs to be remembered so that it is not revisited in vain during the search for targets. Therefore, as proposed in Meghanathan et al. (2019), revisits to distractors are likely made when their locations are forgotten. This implies that there is no difference in VWM processes between first fixations and refixations, since, both involve encoding of distractor locations. Therefore, an attention shift to the subsequent saccade would not cost first fixations and refixations differently, explaining the absence of such a difference in SRPs.

Thus, our finding demonstrates that task-relevance affects oculomotor selection in refixation behavior. Factors affecting saccade planning, which are specific to our task, could explain these results. An interaction between two types of attention is at play in our multitarget visual search task with unrestricted eye movements for 10 s. The first type is top-down attention for detection of task-relevant targets. The second type is oculomotor selection, namely, the attention shift to the next fixation location. Task-relevance of the fixated item biases the competition between strategic and momentary goals of saccade control. Strategic goals in our task involve search for multiple targets and maintenance of already visited targets in memory for the subsequent change detection. In the case of targets, these competing strategic goals introduce a distinction between first fixations and refixations. While first fixations are made as part of a search operation, refixations are made to update items in working memory. Therefore, strategic goals differ between two fixation types. The difference in strategic goals biases momentary goals. Different strategic goals and different working memory processes during first fixations and refixations render disengagement of attention more difficult for first fixations than refixations on targets. Whereas, in the case of distractors, momentary goals dominate in the absence of strategic interest to the location. Since the competition between strategic and momentary goals is different for targets and distractors, we conclude that task-relevance is a factor modulating attention related to saccade planning in natural viewing behavior.

To conclude, task-relevance of an item modulates attentional shifts in preparation for the subsequent saccade in natural viewing behavior. The differences in saccade preparation between first fixations and refixations could be explained by the cost levied on VWM by saccade preparation and by the competition between momentary goals and strategic goals of saccade planning.

## Acknowledgments

The authors were supported by an Odysseus grant (G.0003.12) from the Flemish Organization for Science (FWO) to Cees van Leeuwen.

